# scHiR learns joint cellular representations from paired single-cell transcriptome and 3D genome data

**DOI:** 10.64898/2026.11.06.764816

**Authors:** Yichao Zhao, Pengzhen Jia, Matthew C. Hill, Li Tang, Zhaoyu Fang, Ruiqing Zheng, Min Li

**Author notes:** Correspondence: Min Li,; Ruiqing Zheng.

## Abstract

Simultaneously sequenced scRNA-seq and scHi-C technologies provide matched measurements of gene expression and three-dimensional (3D) genome organization within individual cells, offering complementary insights into cellular state. However, integrative analysis remains challenging due to the heterogeneous structures and characteristics of these modalities. To realize the full potential of such data, we need effective computational methods to learn joint cellular representations. Here, we present scHiR, a framework based on a hyperspherical variational autoen-coder (S-VAE) for learning co-embeddings from paired gene-expression profiles and chromatin contact maps. The framework facilitate downstream analyses such as cell clustering, fine-grained cell-state characterization, and developmental trajectory inference, while modality-specific decoders enable the signal recovery of input profiles. We evaluated scHiR across six paired datasets generated by HiRES, GAGE-seq, and LiMCA. scHiR captured cell-type-specific information and achieved improved clustering performance compared with representative single-omics methods. It was also capable of resolving fine-grained subclusters and revealed discordant and distinct cellular profiles. In developmental biological datasets, scHiR preserved trajectory-related structure and improved pseudotime in agreement with known developmental progression. Finally, scHiR generated reconstructed transcriptomic and chromatin contact profiles with reduced noise, enhanced cell-type-related signals, and improved recovery of held-out chromatin contacts. These results demonstrate that scHiR provides an effective method for learning joint cellular representations from paired scRNA-seq and scHi-C data.

## Introduction

Single-cell sequencing technologies enable the characterization of cellular identity and state. Among these, the transcriptome reflects the functional and regulatory state of a cell, whereas three-dimensional (3D) genome organization reflects the chromatin architecture that provides the spatial context for gene regulation (Bonev and Cavalli 2016). Recent technological advances, such as HiRES, GAGE-seq, and LiMCA, have made it possible to profile the single-cell transcriptome and 3D genome organization simultaneously, generating paired scRNA-seq and scHi-C datasets (Liu et al. 2023; Zhou et al. 2024; Wu et al. 2024). These paired measurements provide an opportunity to investigate the complex relationship between chromatin architecture and gene expression within the same cell.

Although paired scRNA-seq and scHi-C datasets are becoming increasingly available, computational methods specifically designed to learn shared cellular representations from such datasets remain limited. These two measurements of molecular data exhibit differences in noise, scale, and biological meaning. scRNA-seq captures each cell as a gene-expression vector with sparsity and overdispersion. In contrast, scHi-C represents each cell as a high-dimensional chromatin contact matrix, where each value corresponds to a pairwise interaction between two genomic loci. These contact matrices are much sparser and exhibit dependence on genomic distance. In current practice, simultaneously sequenced scRNA-seq and scHi-C data are often analyzed separately using single-omics methods developed for each data type. For scRNA-seq, methods such as scVI, scMGCA, and scGNN learn low-dimensional representations for downstream cell-state analysis (Lopez et al. 2018; Yu et al. 2023; Wang et al. 2021). For scHi-C, methods such as scHiCluster and Fast-Higashi are designed to extract cellular representations from sparse chromatin contact maps (Zhou et al. 2019; Zhang et al. 2022). Although these methods perform well on individual omics datasets, analyzing the two omics separately or directly concatenating their features may fail to capture complementary information in paired data and limit the ability of one data type to help interpret and contextualize the other. Moreover, separately learned latent spaces are not necessarily aligned, making it difficult to obtain a unified representation for individual cells and to model relationships between the two omics effectively.

To address these challenges, we present scHiR, a multi-omics deep framework for learning co-embeddings from simultaneously sequenced scRNA-seq and scHi-C datasets. scHiR uses a hyperspherical variational autoencoder (S-VAE, Davidson et al. 2018)) to integrate gene expression and chromatin interactions into a shared representation for downstream analyses. We demonstrate that scHiR improves cell-type clustering, resolves fine-grained cellular heterogeneity, captures developmental trajectories, and improves joint reconstruction of both omics.

## Results

### scHiR model overview

We designed an S-VAE to jointly extract co-embeddings from simultaneously sequenced scRNA-seq and scHi-C data (Fig. 1). The model takes paired scRNA-seq and scHi-C data from the same cell as input and processes them through two parallel branches. For the scRNA-seq branch, we fed the raw expression values of the top 2,000 highly variable genes (HVGs) into a multilayer perceptron (MLP) encoder to learn transcriptomic representations. For the scHi-C branch, we divided the whole-genome contact map into intra-chromosomal contact matrices for each chromosome. We applied a LeNet-based encoder to each matrix and then averaged the resulting representations to obtain a cell-level scHi-C representation. Next, we concatenated the representations from both omics and used the fused representation to parameterize a von Mises–Fisher (vMF) posterior with mean direction *µ* and concentration parameter *κ*. Following the S-VAE framework, we sampled a shared latent variable *z* from this distribution on the unit hypersphere for reconstruction, providing a common latent representation for each cell.

**Figure 1.**
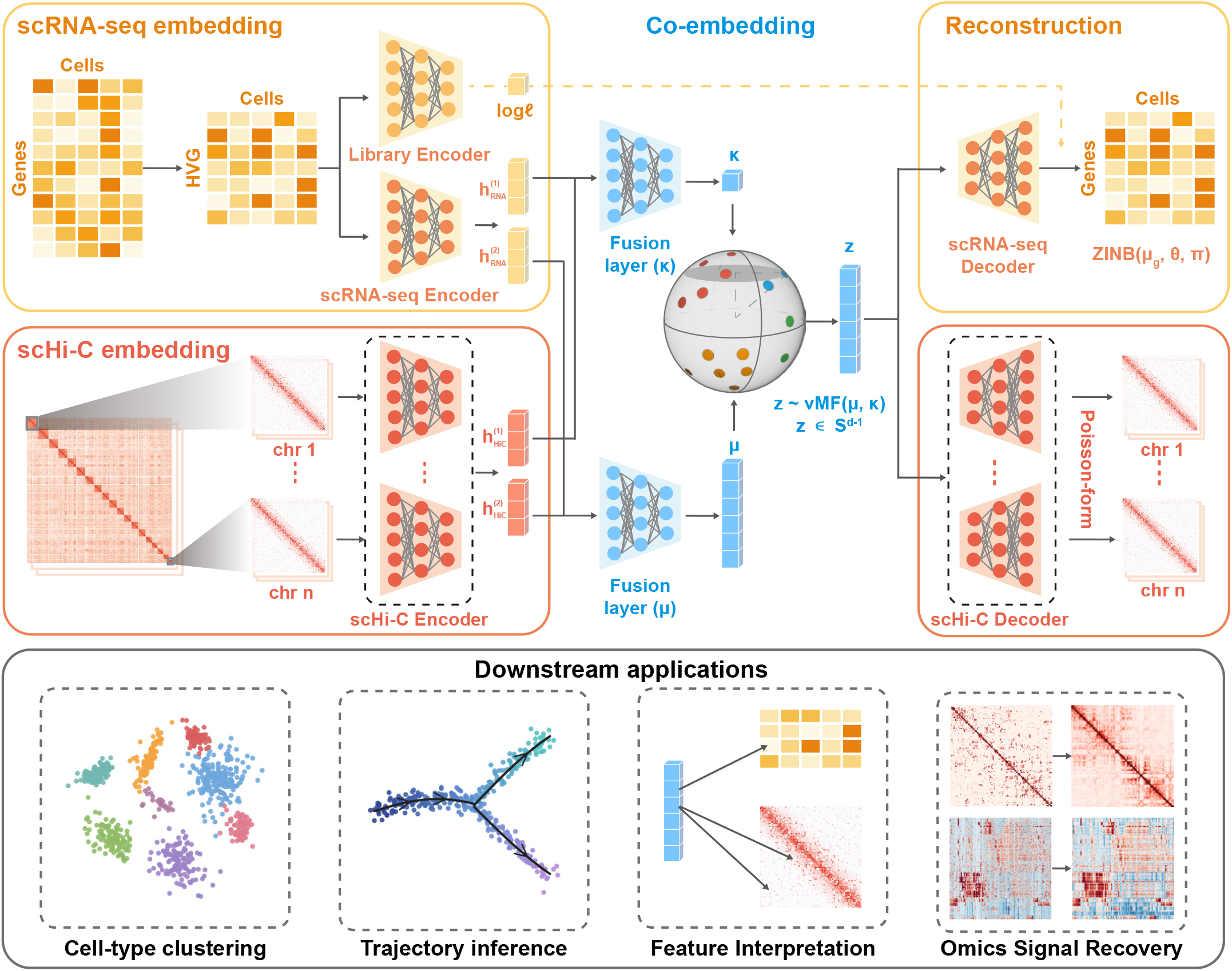
Overview of the scHiR model architecture. Two omics-specific branches process paired scRNA-seq and scHi-C profiles from the same cell. In the scRNA-seq branch, an MLP encoder maps the expression profiles of highly variable genes to a low-dimensional representation. In the scHi-C branch, chromosome-specific LeNet-based convolutional encoders extract features from intra-chromosomal contact matrices, followed by averaging across chromosomes to obtain a cell-level representation. The model fuses the two omics representations and samples a shared latent variable *z* from a von Mises–Fisher distribution, *z* ∼ vMF(*µ, κ*), on the unit hypersphere. Finally, the decoders reconstruct gene expression and chromatin contacts using ZINB and Poisson-based reconstruction objectives, respectively.

The training objective consisted of reconstruction losses and Kullback–Leibler (KL) divergence terms. For scRNA-seq data, we calculated the reconstruction loss using the negative log-likelihood of the ZINB distribution and applied an additional KL divergence term to regularize the inferred library-size latent variable toward an empirical Gaussian prior. For scHi-C data, we used a Poisson-form reconstruction loss on the non-negative contact matrix. We further applied a KL divergence term between the vMF posterior and a uniform prior on the unit hypersphere to regularize the shared latent representation. We also included an anti-collapse regularization term to prevent the hyperspherical mean directions from collapsing to a common direction. After training, we used the joint latent mean direction *µ* as the low-dimensional co-embedding of each cell for clustering, trajectory inference, visualization, and other downstream biological analyses (for further details, see Methods).

To further investigate the biological meaning of the learned co-embeddings, we performed interpretability analysis using the HiRES mouse embryonic stem cell (mESC) dataset by attributing each shared latent dimension to input scRNA-seq genes and scHi-C bin pairs. The resulting attribution profiles differed across latent dimensions and remained relatively balanced between the two omics overall. The latent dimensions were also associated with biological annotations such as cell type and developmental stage. Representative latent dimensions could be further linked to top-attributed genes and scHi-C bin pairs, providing an interpretable view of the biological information encoded by scHiR (Supplemental Fig. S1).

### scHiR effectively learns cell-type-specific latent representations

We first performed clustering analysis using scHiR embeddings on five paired scRNA-seq and scHi-C datasets. These datasets were generated using three different technologies, including HiRES (Liu et al. 2023), GAGE-seq (Zhou et al. 2024), and LiMCA (Wu et al. 2024). Together, they cover a broad range of cell types and sample sizes, including 399 mouse brain cells and 7,469 mouse embryonic stem cells (mESCs) from HiRES, 3,296 mouse cortex cells and 1,445 human bone marrow CD34+ cells from GAGE-seq, and 389 human cell-line cells from LiMCA.

We evaluated the performance by applying k-means clustering to the learned co-embeddings and calculating multiple metrics, including Adjusted Rand Index (ARI), Normalized Mutual Information (NMI), cluster purity, cell-type concentration, and cell-type entropy (see Methods). For the LiMCA cell-line dataset, we used the known cell-line identities as ground-truth labels. For other datasets, we adopted the cell-type annotations provided in the original studies, which were inferred from scRNA-seq analysis and manual curation using known marker genes. For comparison, we included three representative scRNA-seq embedding methods, scVI (Lopez et al. 2018), scMGCA (Yu et al. 2023), and scGNN (Wang et al. 2021), as well as two popular scHi-C embedding methods, FastHigashi (Zhang et al. 2022) and scHiCluster (Zhou et al. 2019).

As shown in Fig. 2A, Supplemental Figs. S2 and S3, scHiR achieved highly competitive performance across all datasets compared with the baseline methods. These results demonstrate that scHiR more effectively learns cell-type-specific latent representations from paired scRNA-seq and scHi-C data than baseline methods that rely on single-omics information alone. For example, in Fig. 2B, scHiR clearly captured the developmental pattern from mesoderm and mesenchymal states to myocytes (Pijuan-Sala et al. 2019) in the mESC dataset, whereas the baseline methods showed less distinct trajectory structure and greater mixing between intermediate cell states.

**Figure 2.**
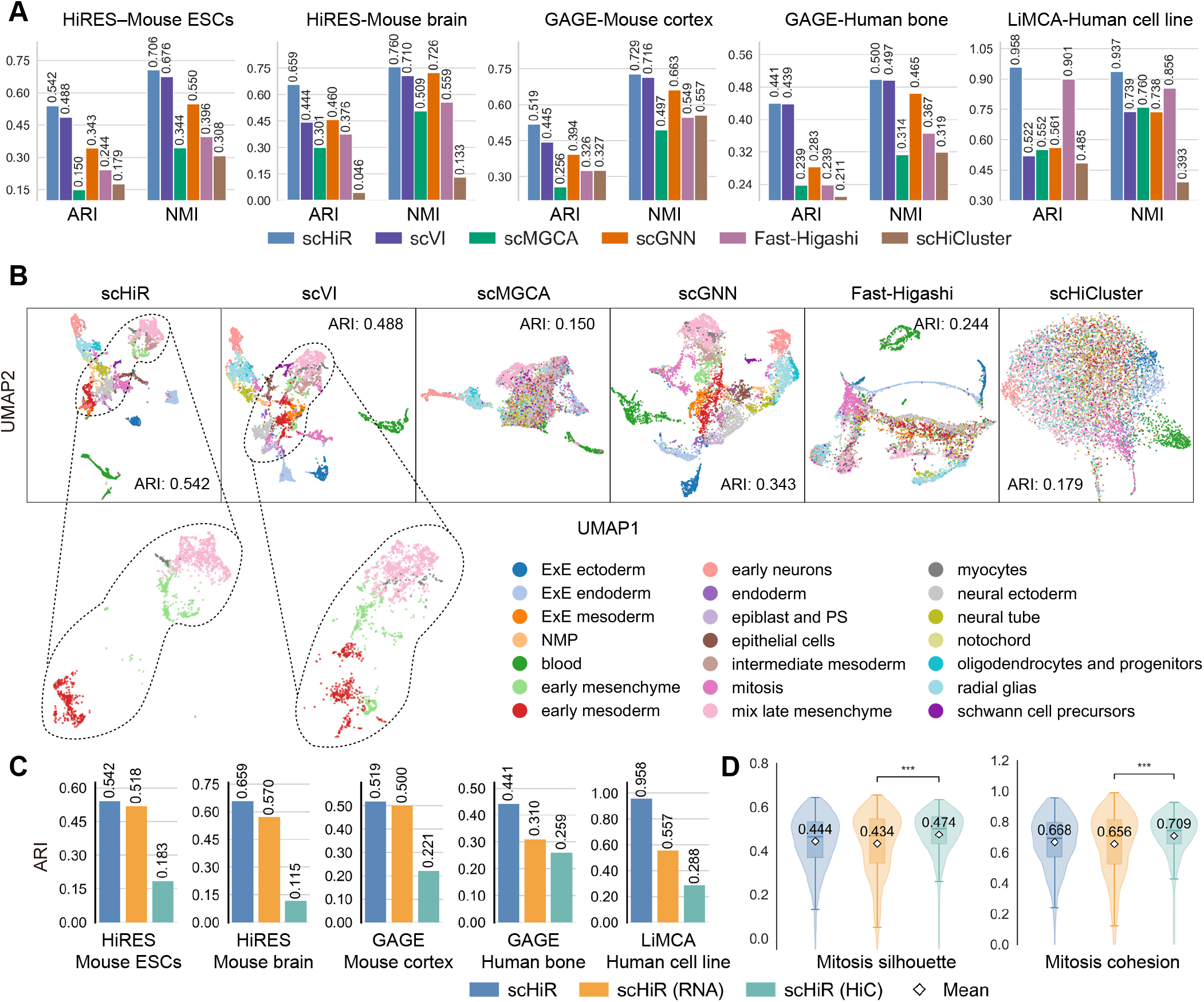
Comparison of clustering performance. (A) Comparison of scHiR with five representative baseline methods across five datasets, evaluated by Adjusted Rand Index (ARI) and Normalized Mutual Information (NMI) after k-means clustering. Full metrics are also provided in Supplemental Table 1. (B) UMAP visualization of the latent representations learned by scHiR and the baseline methods in the HiRES mESC dataset. Each point represents a single cell and is colored according to its annotated cell-type label. (C) Omics ablation analysis comparing ARI between the joint multi-omics scHiR embedding and single-omics scHiR embeddings learned from scRNA-seq alone (scHiR(RNA)) or scHi-C alone (scHiR(HiC)) across five datasets. (D) Comparison of mitosis-cell silhouette score and cohesion across the joint and single-omics scHiR embeddings, where cohesion is defined as the average cosine similarity between mitosis-cell embeddings and their centroid.

To compare the cell-type structure captured by single-omics and multi-omics latent representations, we separately trained scHiR using only scRNA-seq or only scHi-C data, referred to as scHiR(RNA) and scHiR(HiC), respectively. We then compared their embeddings with the joint multi-omics representation. As shown in Fig. 2C and Supplemental Fig. S4A, the joint scHiR representations achieved clustering performance comparable to or better than scHiR(RNA) embeddings and consistently outperformed scHiR(HiC) embeddings. These results indicate that the incorporation of scHi-C information did not weaken the cell-type separation based on transcriptome-based annotations and, in some datasets, further improved clustering performance. For example, in the HiRES mESC dataset, mitosis cells formed a more compact cluster in the scHiR(HiC) embedding than in the scHiR(RNA) embedding. scHiR(HiC) achieved the highest mitosis-cell kNN purity, silhouette score, and cohesion, while the incorporation of scHi-C information enabled the joint scHiR embedding to outperform scHiR(RNA) across these metrics (Fig. 2D and Supplemental Fig. S4B). Together, these results demonstrate that scHiR robustly integrates scRNA-seq and scHi-C information while capturing cell-type-specific structure in the unified latent space.

### scHiR resolves fine-grained cell populations in the mouse brain

In the HiRES mouse brain dataset, we applied Leiden clustering to the scHiR embeddings and identified 12 clusters from the seven original cell-type categories, providing a finer-grained partition of the cells. These clusters largely preserved the original cell-type identities, with only a small number of discordantly annotated cells in clusters C8 and C9 (Fig. 3A,B).

**Figure 3.**
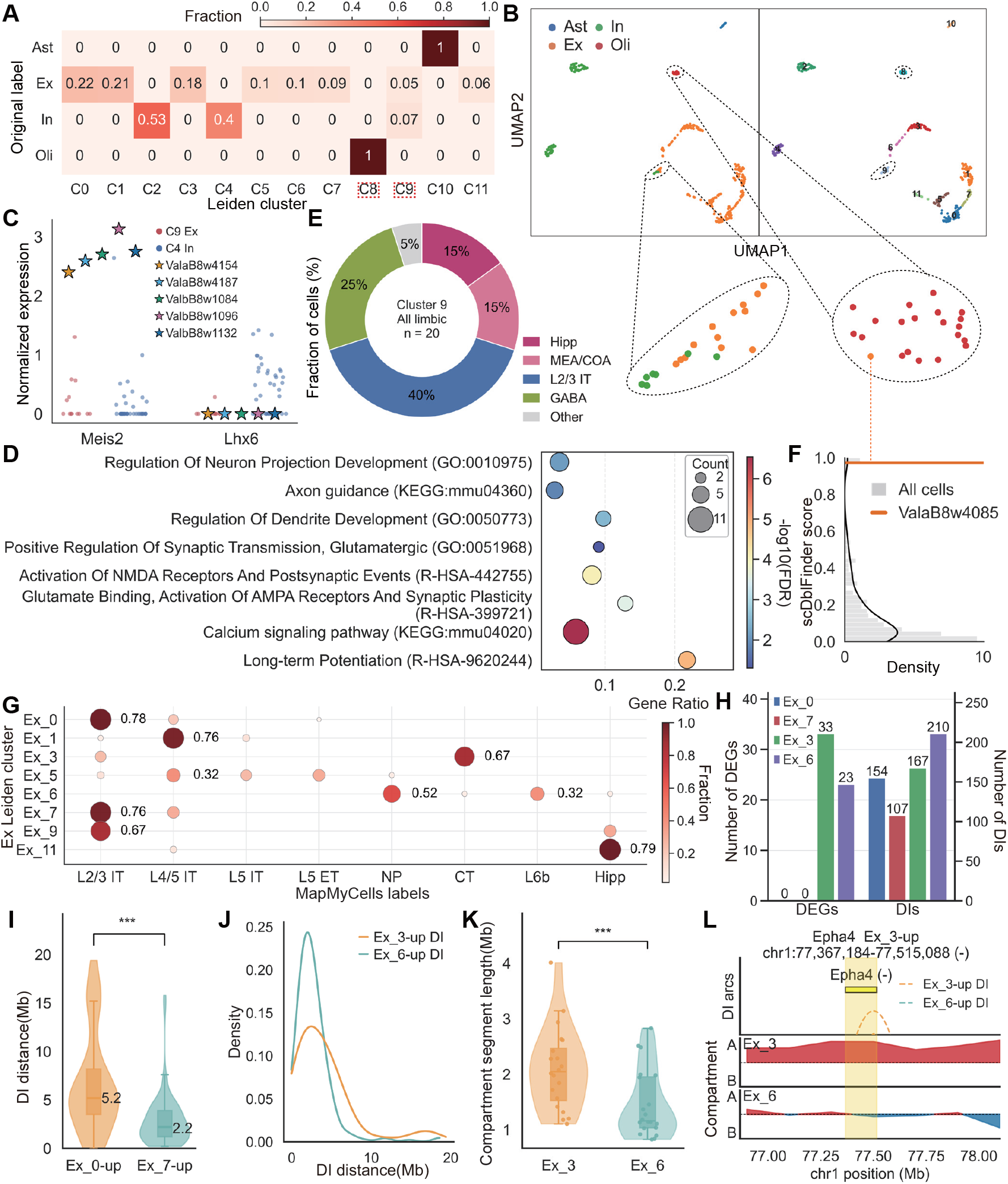
Analysis of fine-grained Leiden clusters in the HiRES mouse brain dataset. (A) Fraction of cells from each original cell-type annotation assigned to each Leiden cluster. (B) UMAP visualization of cells colored by original cell-type annotations (left) and Leiden clustering (right). Dashed circles highlight clusters C8 and C9. (C) Normalized expression of *Meis2* and *Lhx6* in C9-In (n=5), C4-In, and C9-Ex cells. (D) Enrichment analysis of genes commonly up-regulated in both C9-In and C9-Ex relative to C4-In. (E) MapMyCells annotations of all cells in C9. (F) Distribution of scDblFinder doublet scores in cluster C8. The orange line indicates the doublet score of the discordantly annotated cell ValaB8w4085. (G) Composition of Ex Leiden clusters based on MapMyCells cell-type annotations (bootstrap probability *≥* 0.8). (H) Numbers of DEGs and DIs in Ex 0–Ex 7 and Ex 3–Ex 6 comparisons. (I) Distribution of genomic distances for Ex 0-up and Ex 7-up DIs. (J) Density distribution of genomic distances for Ex 3-up and Ex 6-up DIs. (K) Distribution of mean compartment segment lengths per chromosome in Ex 3 and Ex 6. Each dot represents one chromosome. Statistical significance was assessed using a paired Wilcoxon signed-rank test. ns, not significant; ^*^, *P <* 0.05; ^**^, *P <* 0.01; ^***^, *P <* 0.001. (L) Comparison of compartment scores between Ex 3 and Ex 6 around the CT marker gene *Epha4*.

Cluster C9 was predominantly composed of glutamatergic excitatory neurons (Ex) but also contained seven cells annotated as GABAergic inhibitory neurons (In). Reannotation using MapMyCells (RRID:SCR 024672; Supplemental Table 2) reassigned two of them to Ex, whereas the remaining five retained In labels (Supplemental Fig. S5A). We therefore referred to these five cells as C9-In cells (Supplemental Fig. S5B). They expressed In markers while showing limited expression of Ex markers, supporting their inhibitory identity despite clustering with Ex cells (Supplemental Fig. S5C). C9-In cells nevertheless exhibited features distinct from typical GABAergic neurons. Relative to C4-In, they showed increased *Meis2* and decreased *Lhx6* expression (Fig. 3C), consistent with previous findings that *Meis2* promotes GABAergic projection neuron (PN) development whereas *Lhx6* represses this process (Dvoretskova et al. 2024). Moreover, enrichment analysis of commonly up-regulated genes shared between C9-In and C9-Ex revealed pathways related to PN connectivity, glutamatergic signaling, and activity-dependent synaptic plasticity (Fig. 3D). Consistent with these molecular features, MapMyCells assigned all C9 cells to limbic-associated populations (Fig. 3E), with C9-In cells associated with the lateral septum and amygdala-associated nuclei, which contain GABAergic PNs (Anthony et al. 2014; Sun et al. 2019; Keshavarzi et al. 2014). Together, these findings distinguish C9-In from typical inhibitory neurons and suggest the presence of limbic-associated and PN-related features, providing a plausible explanation for their co-clustering with C9-Ex rather than C4-In in the unsupervised embedding.

Cluster C8 contained one cell, ValaB8w4085, that was originally annotated as an Ex but clustered with oligodendrocytes (Oli) in the scHiR embedding. Although MapMy-Cells retained its Ex annotation (Supplemental Fig. S6A), ValaB8w4085 exhibited several Oli-like features. It showed strong expression of Oli marker genes and was consistently more similar to Oli than to Ex cells across all three scHi-C embeddings (Supplemental Fig. S6B,C). To assess whether this discordant profile reflected a potential doublet, we applied scDblFinder (Germain et al. 2022) to its transcriptomic profile, which identified ValaB8w4085 as a putative doublet with a score of 0.97, higher than that of most cells (Fig. 3F). Given its ambiguous cellular identity, we excluded this cell from subsequent cell-type-specific comparisons.

Leiden clustering identified several finer subclusters within the original Ex population. MapMyCells annotation showed that most subclusters corresponded predominantly to specific Ex neuronal subclasses (Fig. 3G). For example, Ex 0, Ex 7, and Ex 9 were predominantly assigned to L2/3 IT neurons, Ex 1 to L4/5 IT neurons, Ex 3 to CT neurons, and Ex 11 to hippocampal neuronal populations (Fig. 3G). Broadly consistent with these annotations, cluster-specific differentially expressed genes (DEGs) showed substantial overlap with subtype marker genes reported by the Allen VISp–ALM transcriptomic taxonomy (Tasic et al. 2018, Supplemental Fig. S7A). According to MapMyCells annotations, Ex 0, Ex 3, Ex 7, and Ex 9 were all associated with L2/3 IT neuronal subclasses but showed distinct region-specific annotation patterns. Ex 0 and Ex 7 were mainly mapped to cortical (CTX) subclasses, Ex 3 was specifically mapped to a retrosplenial cortex (RSP) subclass, and Ex 9 was mapped to piriform/entorhinal cortex (PIR/ENTl) subclasses (Supplemental Fig. S7B). Similarly, Ex 1 and Ex 5 showed distinct L4/5 IT subclass annotation patterns (Supplemental Fig. S7C).

Despite their similar CTX-associated L2/3 IT identities, Ex 0 and Ex 7 formed separate Leiden clusters. We detected no significant DEGs between them but identified hundreds of differential interactions (DIs; Supplemental Table 3), with Ex 0-up DIs spanning significantly longer genomic distances than Ex 7-up DIs (Fig. 3H–I; Supplemental Fig. S8A). We also observed noticeable differences in the relative A/B compartment coverage, while insulation profiles and compartment boundary organization remained similar (Supplemental Fig. S8B–F). Together, these results suggest that Ex 0 and Ex 7 are closely related excitatory neuronal populations with distinct chromatin architecture features. In contrast, Ex 3 and Ex 6 differed at both the transcriptional and chromatin levels (Fig. 3H). Ex 3-up DIs tended to span longer genomic distances (Fig. 3J), whereas Ex 6 contained more compartment boundaries and shorter compartment segments despite broadly similar global A/B composition and insulation profiles (Fig. 3K and Supplemental Fig. S9A–D). We also identified regions showing coordinated transcriptomic and chromatin changes between Ex 3 and Ex 6 (Supplemental Fig. S9E). For example, the CT-associated marker *Epha4* showed higher expression in Ex 3, accompanied by stronger A-compartment features in its surrounding region than in Ex 6 (Fig. 3L).

### scHiR captures cellular developmental trajectories

To investigate whether the latent representations learned by scHiR preserve trajectory-related biological information, we analyzed the mouse main olfactory epithelium (MOE) dataset from LiMCA and the HiRES mESC dataset, both of which have well-established developmental lineages.

We first examined the MOE dataset, which contains 411 cells collected across eight developmental stages. Cells in this dataset progress from progenitor cells to immature olfactory sensory neurons (iOSNs) and mature olfactory sensory neurons (mOSNs). Because the dataset represents a single developmental trajectory with known temporal ordering, it provides a suitable benchmark for quantitative evaluation of trajectory inference. We applied both diffusion pseudotime (DPT, Haghverdi et al. 2016) and Monocle 3 (Cao et al. 2019) to infer pseudotime from the learned embeddings and compared the inferred pseudotime with the developmental stages using Spearman correlation and Kendall’s tau. Using DPT, scHiR embeddings substantially outperformed scHiR(RNA) and scHiR(HiC) embeddings (Fig. 4A), achieving higher Spearman correlation (0.517 versus 0.357 and 0.205) and Kendall’s tau (0.398 versus 0.257 and 0.152). We observed similar improvements using Monocle 3 (Fig. 4B). The pseudotime inferred from scHiR showed a clear correspondence with developmental stages and recovered the expected progenitor–iOSN–mOSN progression (Fig. 4C,D). Partition-based graph abstraction (PAGA, Wolf et al. 2019) analysis further recovered a coherent developmental topology, suggesting a sequential relationship between the two progenitor populations reported in the original study, with progenitor-1 preceding progenitor-2 before differentiation into iOSNs and mOSNs (Fig. 4E and Supplemental Fig. S10).

**Figure 4.**
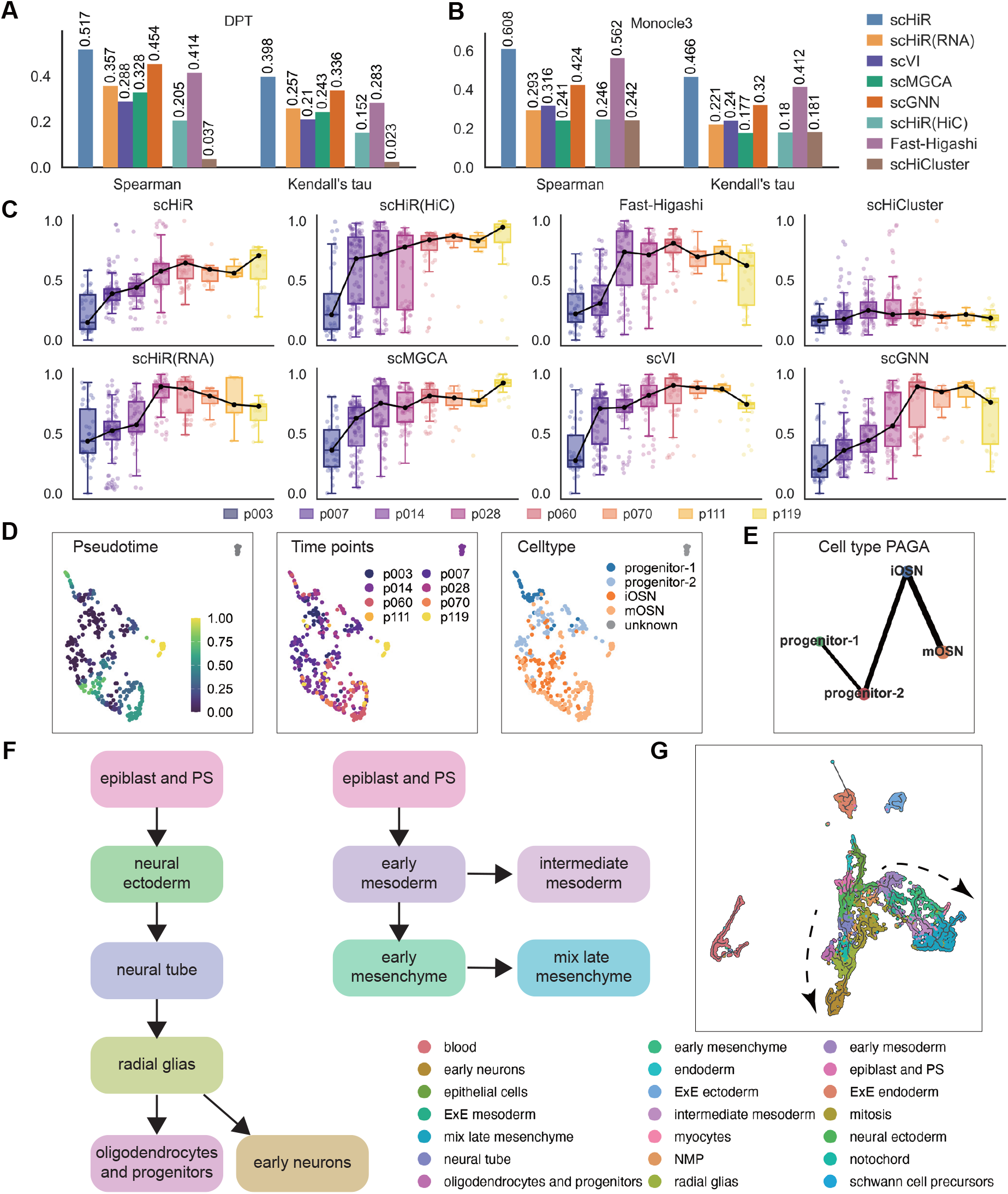
Trajectory analysis of the LiMCA MOE and HiRES mESC datasets. (A,B) Spearman correlation and Kendall’s tau between developmental stages and pseudotime inferred by DPT (A) or Monocle 3 (B) on the MOE dataset. For DPT, the reported values are the mean correlation coefficients across all valid p003 cells used as root cells. (C) Distribution of DPT pseudotime across developmental stages for each embedding method on the MOE dataset using the root cell with the highest Spearman correlation. Black lines indicate stage-wise median pseudotime. (D) UMAP plot of Monocle 3 trajectory inference from the scHiR embedding on the MOE dataset. Cells are colored by normalized pseudotime (left), developmental stages (middle), and annotated cell types (right). (E) PAGA graphs constructed for the scHiR embedding on the MOE dataset using annotated cell types as groups. Edge thickness indicates PAGA connectivity strength, and weak edges were omitted for visualization. (F) Schematic summary of the two major trajectories represented in the mESC dataset. (G) UMAP plot of Monocle 3 trajectory inference from the scHiR embedding in the mESC dataset. Cells are colored by cell types.

We next evaluated scHiR on the HiRES mESC dataset, which contains a more complex developmental process involving multiple branching lineages. The scHiR embedding retained developmental-stage information, with cells from successive stages forming a continuous trajectory in the latent space (Supplemental Fig. S11A,B). We further applied Monocle 3 and PAGA to investigate lineage relationships among embryonic cell types. Consistent with the original study, the scHiR embedding recovered the early neuron trajectory and the mixed late mesenchyme trajectory (Fig. 4F,G and Supplemental Fig. S11C). Collectively, these results suggest that integrating transcriptomic and chromatin contact information allows scHiR to capture developmental stage-associated variation in its latent representations, providing a useful basis for downstream pseudotime analysis.

### scHiR improves transcriptomic and chromatin reconstruction through multi-omics learning

In GAGE-seq mouse cortex cells, we next evaluated how joint learning of transcriptomic and chromatin contact information affects reconstruction across both omics.

For the transcriptomic output, we first compared reconstructed HVG expression profiles with the input scRNA-seq profiles. At the gene level, scHiR achieved significantly higher correlations with the input expression profiles than the separately trained RNA-only model scHiR(RNA) (median Pearson correlation = 0.193 vs. 0.165, paired Wilcoxon test *P* = 8.899 *×* 10^−157^; Fig. 5A). Consistently, marker-gene analysis showed that scHiR recovered a larger fraction of input-derived cell-type marker genes than scHiR(RNA)

**Figure 5.**
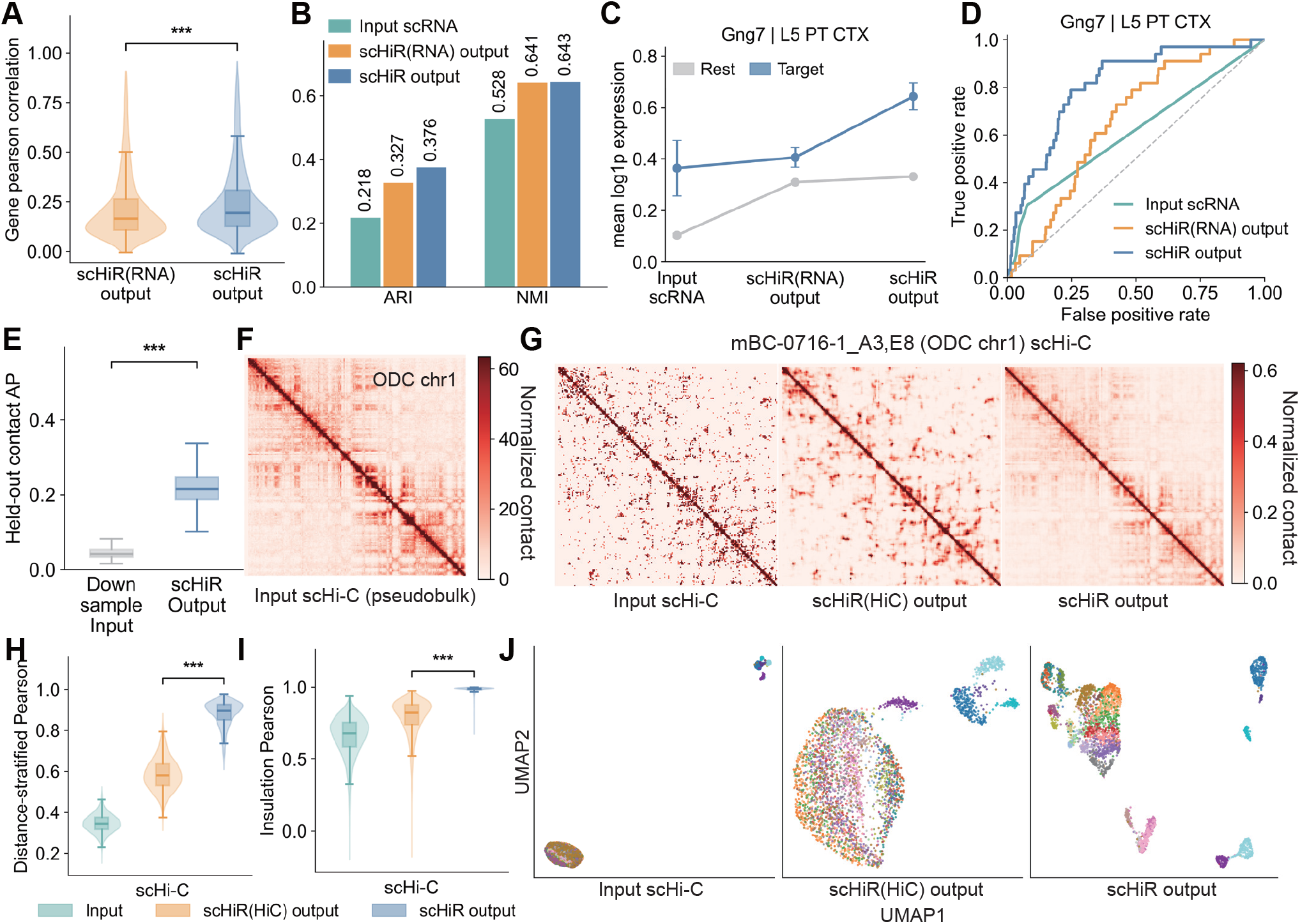
Multi-omics learning in scHiR improves scRNA-seq and scHi-C reconstruction in GAGE-seq mouse cortex cells. (A) Gene-wise Pearson correlation between reconstructed and input scRNA-seq profiles for scHiR(RNA) output and scHiR output. (B) Cell-type clustering performance of scRNA-seq-derived representations from input scRNA-seq, scHiR(RNA) output, and scHiR output, evaluated by ARI and NMI using annotated cell types as ground truth. (C) Mean expression of the L5 PT CTX marker gene *Gng7* in target and non-target cells across input scRNA-seq, scHiR(RNA) output, and scHiR output. (D) ROC curves showing the ability of *Gng7* expression to distinguish L5 PT CTX cells from other cells across the three scRNA-seq profiles. (E) Average precision (AP) for recovering held-out contacts from downsampled input scHi-C contact maps using scHiR. (F) Input pseudobulk scHi-C contact map of chromosome 1 in oligodendrocytes (ODCs). (G) Representative ODC chromosome 1 scHi-C contact maps from input scHi-C, scHiR(HiC) output, and scHiR output. (H) Distance-stratified Pearson correlations between each contact map and the input pseudobulk scHi-C reference matched by cell type. (I) Insulation Pearson correlations between each contact map and the input pseudobulk scHi-C references matched by cell type. (J) UMAP visualization of scHi-C-derived representations from input scHi-C, scHiR(HiC) output, and scHiR output. Cell-type colors are consistent with those used in Supplemental Fig. S3B. Statistical significance was assessed using a paired Wilcoxon signed-rank test. ns, not significant; ^*^, *P <* 0.05; ^**^, *P <* 0.01; ^***^, *P <* 0.001.

(Supplemental Fig. S12A,B). k-means clustering of reconstructed HVG expression profiles further showed that incorporating scHi-C information achieved comparable performance to scHiR(RNA) when using annotated cell types as ground truth (Fig. 5B), and in some cases further improved performance. For example, in scHiR output, the L5 PT CTX marker *Gng7* showed a greater expression elevation between L5 PT CTX cells and the remaining cell types than in the input scRNA-seq profiles or scHiR(RNA) output (Fig. 5C,D), indicating that cell-type-specific transcriptomic information is enhanced by the scHiR model. Similarly, we observed comparable improvements for several other marker genes across different cell types (Supplemental Fig. S12C,D).

We next examined whether scHiR improves the reconstruction of sparse scHi-C contact maps and helps generate denoised chromatin contact patterns. We first downsampled the observed counts in each input scHi-C contact map to 30% of the original depth using binomial thinning and used them as input. In the reconstructed outputs, scHiR recovered held-out contacts with higher average precision (AP) and area under the receiver operating characteristic curve (AUROC) than the downsampled input baseline (Fig. 5E and Supplemental Fig. S13A,B), supporting its ability to partially recover missing chromatin contacts from sparse input scHi-C data. We then evaluated reconstructed scHi-C contact maps for their ability to capture global cell-type-level patterns by comparing them with cell-type-matched input pseudobulk scHi-C references. Visual comparison of representative ODC chromosome 1 contact maps showed that scHiR output most closely resembled the input pseudobulk scHi-C reference (Fig. 5F,G and Supplemental Fig. S13C). Consistent with this observation, scHiR output also showed higher distance-stratified and insulation Pearson correlation with the input pseudobulk scHi-C references than both input scHi-C and scHiR(HiC) output (Fig. 5H,I).

Finally, k-means clustering of scHi-C-derived representations showed that scHiR output achieved substantially higher cell-type clustering accuracy than both input scHi-C and scHiR(HiC) output (ARI = 0.426 vs. 0.141 and 0.173; NMI = 0.699 vs. 0.413 and 0.444; Supplemental Fig. S13D), while UMAP visualization further showed clearer cell-type separation in scHiR output (Fig. 5J), suggesting that the incorporation of transcriptomic information enables scHiR to learn more cell-type-specific chromatin contact patterns than relying on scHi-C data alone.

## Discussion

Simultaneously sequenced scRNA-seq and scHi-C technologies provide an opportunity to examine transcriptomic state and 3D genome organization within individual cells. However, the two omics differ substantially, making it challenging to directly learn an integrated representation, limitating the development of computational methods for exploring information from such datasets. In this study, we developed scHiR, a multi-omics deep learning framework that combines omics-specific encoders with a shared S-VAE latent space to learn co-embeddings from paired scRNA-seq and scHi-C datasets. By jointly optimizing reconstruction losses together with latent-space regularization, scHiR learns a unified cellular representation that integrates information from both omics.

Across five paired scRNA-seq and scHi-C datasets generated by different technologies, scHiR consistently captured cell-type-specific information and achieved competitive or improved clustering performance compared with representative single-omics methods. Importantly, the joint embedding performed comparably to or better than embeddings trained only with scRNA-seq data while clearly outperforming embeddings trained only with scHi-C data in most settings, indicating that the integration of chromatin contact maps provided complementary information in specific biological contexts.

The integrated representation learned by scHiR also enabled fine-grained biological analyses beyond broad cell-type classification. In the HiRES mouse brain dataset, Leiden clustering of scHiR embeddings resolved neuronal subclusters that were not fully captured by the original annotations. Some clusters were associated with distinct transcriptomic patterns, while others showed more prominent differences in chromatin organization. In addition, we revealed cells with discordant annotations or distinct profiles, including inhibitory neurons that clustered with excitatory neurons and a putative doublet. These observations suggest that learned co-embeddings can help refine cell-state annotations and prioritize cell populations for detailed downstream characterization.

Developmental analyses further showed that scHiR preserved trajectory-related information in the shared latent space. In the LiMCA MOE dataset, scHiR recovered the expected progression from progenitor cells to immature and mature olfactory sensory neurons and improved pseudotime agreement with developmental stages compared with single-omics embeddings. In the HiRES mESC dataset, scHiR also captured continuous developmental organization and recovered major lineage relationships. Collectively, given that changes in gene expression and 3D chromatin organization may not occur simultaneously during cellular differentiation, scHiR has the potential to better characterize gradual cell-state transitions by integrating complementary information from both omics.

In addition to representation learning, scHiR supported the reconstruction and signal recovery of both transcriptomic and chromatin contact profiles through joint multi-omics training. For scRNA-seq, compared with the input profiles, the reconstructed expression profiles showed enhanced cell-type information and clearer marker-gene signals. For scHi-C, scHiR recovered partial held-out contacts from downsampled sparse contact maps, and the incorporation of transcriptomic information helped produce reconstructed contact maps that more closely matched cell-type-level pseudobulk references. These results indicate that paired learning can provide cross-omics context for reducing sparsity-associated noise and generating reconstructed profiles with enhanced cell-type information.

Despite these strengths, several limitations of the current work should be acknowledged. scHiR currently focuses on intra-chromosomal contact matrices at a fixed resolution, which may limit its ability to model finer-scale chromatin contacts or interchromosomal interactions. Moreover, although scHiR provides interpretable latent dimensions and enables paired downstream analyses, additional experimental evidence would be needed to establish regulatory relationships between chromatin architecture and gene expression. Future extensions could incorporate multi-resolution chromatin features, batch-aware integration across technologies, and model-based perturbation or mechanistic analyses to further investigate regulatory relationships between chromatin architecture and gene expression.

## Methods

### Data preprocessing

We collected publicly available simultaneously sequenced scRNA-seq and scHi-C datasets from the Gene Expression Omnibus (GEO). We obtained the mouse brain and mouse embryonic stem cell (mESC) datasets from HiRES, which include 399 mouse brain cells and 7,469 mESCs (GEO: GSE223917, Liu et al. 2023). We also collected 389 human cellline cells and 411 main olfactory epithelium (MOE) cells from LiMCA (GEO: GSE240128, Wu et al. 2024). Additionally, we obtained the mouse cortex and human bone marrow CD34+ datasets from GAGE-seq, which include 3,296 and 1,445 cells, respectively (GEO: GSE238001, Zhou et al. 2024).

For scRNA-seq data, we first removed cells with no detected gene expression and used Scanpy (Wolf et al. 2018) to identify the top 2,000 highly variable genes (HVGs) after library-size normalization and log-transformation. To retain count-valued inputs for the zero-inflated negative binomial (ZINB) reconstruction model, we used the raw count matrix of these selected HVGs, rather than normalized expression values as the model input.

For scHi-C data, we processed each chromosome matrix independently. We first divided the whole-genome contact matrix of each cell into intra-chromosomal contact matrices for each chromosome at 1 Mb resolution. To reduce the strong right skew of raw contact distributions and stabilize variance, we applied a log transformation to each contact matrix. We then truncated extreme values at the 99th percentile of each matrix to reduce the influence of outliers. Finally, we scaled each clipped matrix by its maximum value to normalize the input range to approximately [0,1], as shown in Equation 1:

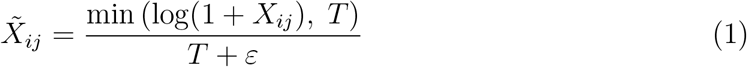

where *X*_*ij*_ denotes the raw contact count between genomic bins *i* and *j, T* is defined as the 99th percentile of the log-transformed contact matrix, and *ε* is a small constant added to avoid division by zero.

### Model architecture

We designed a hyperspherical variational autoencoder (S-VAE, Davidson et al. 2018) to jointly learn low-dimensional co-embeddings from simultaneously measured scRNA-seq and scHi-C data from the same cells. The model consisted of two omics-specific encoder branches, a shared hyperspherical latent space, and two omics-specific decoders for reconstructing the input data.

For the scRNA-seq encoder, we used a multilayer perceptron (MLP) to progressively reduce the dimensionality of the gene expression profile. We then used three additional fully connected blocks to refine high-level transcriptomic features. Each fully connected block consisted of a linear transformation with 128 hidden units, batch normalization, LeakyReLU activation, and dropout regularization. Finally, the encoder mapped the hidden representation to two 128-dimensional branch-specific feature vectors, denoted as 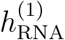 and 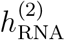.

For scHi-C, we used an independent LeNet-based convolutional encoder for each chromosome. Each encoder processed an intra-chromosomal contact matrix through two 2D convolutional layers followed by average pooling to capture local chromatin interaction patterns and reduce the matrix dimensions. We then flattened the resulting feature maps and passed them through a fully connected layer to generate chromosome-level hidden representations. Two separate linear layers further mapped each chromosome-level representation to two 128-dimensional feature vectors. We averaged these feature vectors across chromosomes to obtain two cell-level scHi-C feature vectors, denoted as 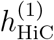 and 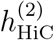.

We fused the two omics by concatenating 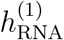 with 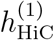 and 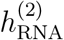 with 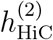, and passed each concatenated feature through a linear fusion layer followed by a final linear projection layer. This process produced two raw vMF parameter outputs: a *d*-dimensional vector *h*^(1)^ for computing the mean direction and a scalar *h*^(2)^ for computing the concentration parameter. We then obtained the vMF parameters as follows:

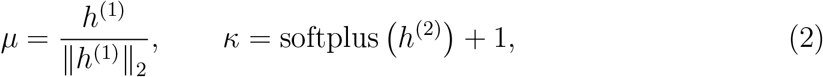

where *µ* denotes the mean direction on the unit hypersphere, and *κ* controls the concentration of the posterior distribution around *µ*. We then sampled the shared latent variable from a vMF posterior:

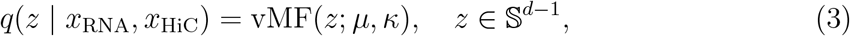

where *d* is the dimension of the shared hyperspherical latent space, *q*(*z* | *x*_RNA_, *x*_HiC_) denotes the shared posterior distribution, and *z* is the sampled latent variable on the unit hypersphere.

We used the uniform distribution on the unit hypersphere, *p*(*z*) = Unif(*S*^d−1^), as the prior and regularized the vMF posterior toward this prior using the Kullback–Leibler (KL) divergence, *D*_KL_(*q*(*z* | *x*_RNA_, *x*_HiC_) ∥ *p*(*z*)).

The decoders reconstructed transcriptomic and scHi-C profiles from the shared latent variable *z*. For scRNA-seq data, we modeled gene expression with a zero-inflated negative binomial (ZINB) distribution to account for sparsity and overdispersion (Lopez et al. 2018). The scRNA-seq decoder estimated the ZINB parameters, including the zero-inflation probability *π*, mean *µ*_*g*_, and dispersion parameter *θ*.

To correct for sequencing-depth differences across cells, we used a separate library-size encoder to infer a cell-specific latent library size factor. The scRNA-seq decoder first generated gene-wise normalized expression proportions *ρ*, and then scaled these proportions by the inferred library size to obtain the final mean parameter:

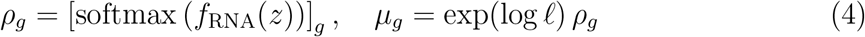

where *l* denotes the inferred cell-specific library size, *ρ*_*g*_ denotes the decoder-predicted expression proportion for gene *g*, and *f*_RNA_(·) denotes the scRNA-seq decoder network. For scHi-C reconstruction, we used chromosome-specific transposed-convolution decoders corresponding to the LeNet-based encoders. For each chromosome, the decoder mapped the shared latent variable *z* back to the intra-chromosomal contact space and produced an unconstrained contact score matrix. We then transformed the score matrix into non-negative contact-intensity parameters using a softplus function:

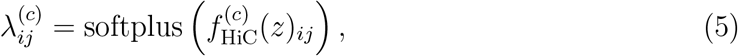

Where 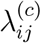 denotes the reconstructed non-negative contact-intensity parameter for the preprocessed contact value between genomic bins *i* and *j* on chromosome *c*, and 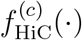 denotes the chromosome-specific scHi-C decoder. We defined the scHi-C reconstruction loss as a Poisson-form reconstruction loss implemented with torch.nn.functional.poisson nll loss, using the preprocessed input contact-intensity matrix as the reconstruction target.

The training objective combined the scaled scRNA-seq reconstruction and librarysize regularization loss, the weighted scHi-C Poisson-form reconstruction loss, the vMF- to-uniform KL divergence for the shared latent variable, and a batch-level anti-collapse regularization term:

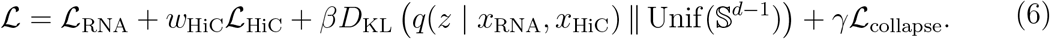

Here, *ℒ*_RNA_ includes the ZINB negative log-likelihood and the KL regularization term for the inferred library-size latent variable, scaled by 1000*/G*, where *G* is the number of selected HVGs. The parameter *w*_HiC_ balances the contribution of the scHi-C recon-struction loss and was automatically adjusted after an initial warm-up period according to the relative scales of the scRNA-seq and scHi-C losses. The parameter *β* controls the strength of the vMF-to-uniform KL regularization and was linearly increased to a maximum value of 0.05 during training. The parameter *γ* controls a batch-level anti-collapse regularization term that discourages excessive alignment of cell-wise mean directions:

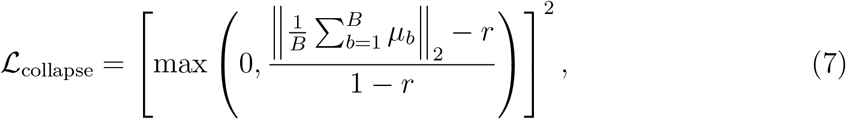

where *B* denotes the batch size, *µ*_*b*_ denotes the unit mean direction of the *b*-th cell in the batch, and *r* denotes a tolerance threshold. This term remains inactive when the batch-averaged direction is sufficiently dispersed.

scHiR was trained using the Adam optimizer with an initial learning rate of 5 *×* 10^−4^. We set the dimension of the shared hyperspherical latent space to 32, and the batch size to 32 for all datasets. To ensure sufficient training across datasets with different sample sizes, we trained scHiR for 200 epochs on the HiRES mESC and GAGE-seq mouse cortex datasets, which contained substantially more cells, and for 100 epochs on the remaining smaller datasets. After training, we used the joint latent mean direction *µ* as the low-dimensional co-embedding of each cell for clustering, trajectory inference, visualization, and downstream biological interpretation.

To compare the representations learned from individual omics types, we trained two single-omics versions of scHiR using the same hyperspherical latent framework. For scHiR(RNA), we retained only the scRNA-seq encoder, the shared vMF latent layer, and the scRNA-seq decoder, and optimized the ZINB reconstruction loss together with the library-size regularization and vMF-to-uniform KL regularization. For scHiR(HiC), we retained only the chromosome-specific scHi-C encoders, the shared vMF latent layer, and the chromosome-specific scHi-C decoders, and optimized the scHi-C Poisson-form reconstruction loss together with the vMF-to-uniform KL regularization. In both versions, we removed the unavailable omics-specific branch and its reconstruction loss while retaining the same shared hyperspherical latent dimension and training settings as in the full model.

### Clustering analysis

For baseline methods, including scVI, scMGCA, scGNN, Fast-Higashi, and scHiCluster, we followed the preprocessing and parameter settings recommended in the original publications or official implementations. For each embedding, we applied k-means clustering using scikit-learn (Pedregosa et al. 2011). We used 20 random initializations and set the number of clusters equal to the number of annotated cell types. We then evaluated clustering performance using multiple complementary metrics by comparing predicted k-means clusters with annotated cell-type labels:

Adjusted Rand Index (ARI) measures whether pairs of cells are consistently assigned in the annotated cell-type labels and predicted k-means clusters.

Normalized Mutual Information (NMI) measures the shared information between predicted k-means clusters and annotated cell-type labels.

Let *M* ∈ ℝ^*C×K*^ denote the contingency matrix between annotated cell types *C* and predicted k-means clusters *K*, where *M*_*ij*_ represents the number of cells from cell type *i* assigned to cluster *j*. We row-normalized *M* to obtain *M* ^row^ and column-normalized *M* to obtain *M* ^col^. We define cluster purity, which measures the average dominance of the major cell type within each predicted k-means cluster, as:

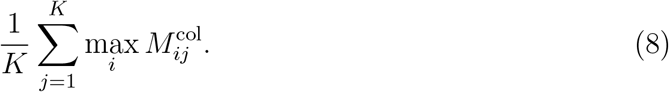

We define cell-type concentration, which measures how strongly cells from the same annotated cell type are assigned to the same predicted k-means cluster, as:

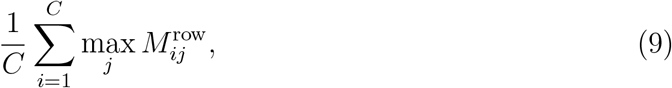

We define cell-type entropy, which measures the degree of mixing of cells from the same annotated cell type across predicted k-means clusters, as:

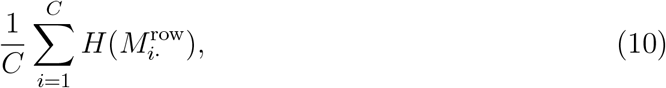

where *H*(·) denotes Shannon entropy. Lower entropy indicates better clustering consistency for each annotated cell type.

To further compare the local organization of mitosis cells between the single- and multi-omics embeddings in the HiRES mESC dataset, we calculated mitosis-cell k-nearest neighbor (kNN) purity, silhouette score, and cohesion. kNN purity measures the fraction of mitosis cells among the nearest neighbors of each mitosis cell in the embedding space. The score ranges from 0 to 1, where higher values indicate stronger local neighborhood enrichment of mitosis cells (Huizing et al. 2023). The silhouette score quantifies how well mitosis cells separate from other cells based on cosine distances in the embedding space, with higher values indicating better separation. For each cell *i*, we calculated the silhouette score as

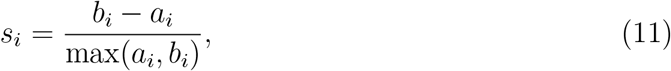

where *a*_*i*_ denotes the mean distance from cell *i* to other cells in the target population, and *b*_*i*_ denotes the mean distance from cell *i* to cells outside the target population. We averaged *s*_*i*_ over mitosis cells. Cohesion measures the compactness of the mitosis cell population. For the target cell population *C*, we calculated cohesion as the average cosine similarity between each cell embedding and the normalized centroid of that population:

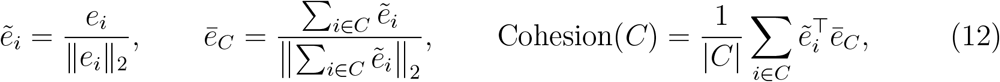

where *e*_*i*_ denotes the embedding of cell *i*, 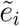 denotes its L2-normalized embedding, 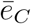 denotes the normalized centroid direction of population *C*, and |*C*| denotes the number of cells in population *C*. Higher values indicate a more compact cell population in the latent space.

### Cell subtype annotation and refinement

For fine-grained subtype analysis in the HiRES mouse brain dataset, we performed Leiden clustering on the 32-dimensional scHiR latent embeddings. We constructed a k-nearest-neighbor graph and performed Leiden clustering using Scanpy with default parameters (n neighbors = 15, resolution = 1) and assigned each Leiden cluster to the original cell-type annotation. To further characterize the cell identities of Leiden clusters, we annotated cells using the HierarchicalMap workflow of MapMyCells (RRID:SCR 024672) with the Allen Brain Cell Atlas 10x Whole Mouse Brain reference taxonomy (CCN20230722, Yao et al. 2023). HierarchicalMap assigns cells across multiple levels of the Allen taxonomy and provides confidence estimates through bootstrap probabilities. To evaluate whether discordantly annotated cells represented potential doublets, we applied scD-blFinder (Germain et al. 2022) to the raw scRNA-seq UMI count matrix from the HiRES mouse brain dataset. For Ex-subcluster visualization, we retained cells with subclass bootstrap probabilities *≥* 0.8 and excluded subclasses whose highest fraction in any Ex cluster was below 0.15. For detailed visualization of L2/3 IT- and L4/5 IT-associated Ex clusters, we used MapMyCells cluster-level annotations and retained labels represented by at least three cells.

### Differential expression analysis

For differential expression analysis, we first normalized raw scRNA-seq data to a total count of 10^4^ per cell and log-transformed using Scanpy (Wolf et al. 2018). We performed differential expression analysis using the Wilcoxon rank-sum test implemented in `scanpy.tl.rank_genes_groups`, and then corrected *P* values for multiple testing using the Benjamini–Hochberg procedure to control the false discovery rate (FDR).

We defined significant genes as those with an *FDR <* 0.05 and an absolute log-fold change (|logFC|) *>* 0.5 for most analyses. However, because we only identified five discordant cells (C9-In) in cluster C9, we used more relaxed thresholds for DEG identification in this comparison. We retained genes showing consistent directional changes in C9-In and C9-Ex cells relative to the C4-In population, with an absolute difference in mean log-normalized expression of at least 0.25 and a nominal *P <* 0.05.

For functional enrichment analysis, we used Enrichr through the GSEApy package (Fang et al. 2023). We analyzed DEGs using Gene Ontology Biological Process, KEGG, and Reactome gene-set collections and considered terms with an adjusted *P <* 0.05 to be significantly enriched.

### Chromatin architecture analyses

We used DISHIC (Zhao et al. 2025) to identify DIs from scHi-C data. We first converted intra-chromosomal contacts into 200-kb genomic bin pairs and ran DISHIC separately for each chromosome using 200-kb genomic feature files to obtain raw *P* values. To reduce sparsity-driven signals, we retained cis interactions within 20 Mb that were supported in either comparison group by at least 10 cells. For the retained interactions, we adjusted *P* values within each chromosome and genomic-distance group using the Benjamini– Hochberg procedure. Significant DIs were defined as interactions with an FDR *<* 0.1 and an absolute interaction logFC *>* 0.5.

For A/B compartment analysis, we pooled raw scHi-C contact pairs from cells in each group to generate pseudobulk scHi-C at 200-kb resolution. We retained cis contacts and created balanced cooler files using cooler (Abdennur and Mirny 2020). We then calculated the first compartment eigenvector (E1) using `cooltools.eigs_cis` with GC content as the phasing track. Bins with positive E1 values were assigned to the A compartment, whereas bins with negative E1 values were assigned to the B compartment. We defined compartment boundaries as changes in compartment state (A-to-B or B-to-A) between adjacent genomic bins. For each chromosome, we calculated the number of compartment boundaries per 100 Mb and the mean length of continuous A/B compartment segments. We also calculated the total genomic lengths of A and B compartments by summing the lengths of bins assigned to each compartment. To assess compartmental interaction preferences, we quantified the fractions of A-A, B-B, and A-B interactions using balanced contact counts.

For insulation analysis, we calculated genome-wide insulation scores from 200-kb balanced pseudobulk cooler files using `cooltools.insulation` with a 1-Mb window, ignoring the first diagonal. We used the resulting log_2_ insulation scores for downstream analyses. To compare insulation profiles, we calculated Pearson and Spearman correlations between insulation scores from matching genomic bins.

### Trajectory analysis

For trajectory topology analysis, we constructed a kNN graph from ceh embedding using Scanpy (sc.pp.neighbors). We then performed PAGA using either developmental stages or annotated cell types as grouping labels. To infer pseudotime, we computed diffusion maps using Scanpy (`sc.tl.diffmap`) based on the kNN graph and subsequently inferred DPT using `sc.tl.dpt`. We also projected embeddings into a two-dimensional UMAP space and learned trajectory graphs using Monocle 3.

Depending on the dataset and trajectory method, we selected root cells according to the original annotations. In the MOE dataset with a single developmental trajectory, we used all cells from the earliest stage (p003) as candidate root cells. For DPT, we ran pseudotime inference separately using each p003 cell as the root and reported the mean Spearman correlation and Kendall’s tau across all valid roots. For Monocle 3 trajectory inference, we selected all p003 cells as root cells. In the mESC dataset, we selected all cells annotated as epiblast and primitive streak (epiblast and PS) as root cells for cell-type trajectory inference.

### Reconstruction evaluation

We evaluated whether the reconstructed scRNA-seq and scHi-C profiles preserved cell-type structure through clustering analysis. For scRNA-seq, we first transformed the input HVG expression profiles, scHiR(RNA) outputs, and scHiR outputs by log(1 + *x*), standardized them with zero mean and unit variance, and reduced them to 30 principal components by PCA. For scHi-C, we retained upper-triangular entries at genomic distances of 1–30 Mb within each chromosome and concatenated the corresponding contact values across chromosomes for each cell. We then standardized and reduced them to 30 principal components by PCA. Using the resulting scRNA-seq- and scHi-C-derived representations, we performed k-means clustering and evaluated the performance using adjusted Rand index (ARI) and normalized mutual information (NMI) against annotated cell-type labels.

To investigate the ability of scHiR to recover missing chromatin interactions rather than simply smooth observed contacts in sparse scHi-C maps, we performed a downsampling analysis. For each input scHi-C contact map, we downsampled the observed contact counts to 30% of the original depth using binomial thinning. We defined positions with nonzero counts in the full input map but zero counts after downsampling as positive held-out contacts, and positions with zero counts in both the full and downsampled maps as negatives. We then used reconstructed scHi-C contact values as prediction scores and evaluated held-out contact recovery using average precision (AP) and area under the receiver operating characteristic curve (AUROC). Here, we treated the downsampled input map as a no-recovery baseline because all candidate positions had zero values.

To evaluate whether reconstructed scHi-C contact maps captured global cell-type-level chromatin contact patterns, we generated input pseudobulk scHi-C references by summing the corresponding input scHi-C contact maps for each annotated cell type. When comparing a scHi-C contact map with pseudobulk scHi-C, we used a leave-one-out strategy and excluded the target cell when constructing its matched pseudobulk reference. We then calculated similarity using distance-stratified Pearson correlation and insulation Pearson correlation. For distance-stratified Pearson correlation, we grouped contacts into genomic distance strata of 1–2, 3–5, 6–10, 11–20, and 21–50 Mb, computed Pearson correlation within each stratum, and averaged the resulting correlations across strata. For insulation Pearson correlation, we computed insulation profiles using a 5-bin window.

## Data Access

All datasets analyzed in this study are publicly available from the Gene Expression Omnibus (GEO). The HiRES mouse brain and mESC datasets are available under accession GSE223917; the LiMCA human cell-line and mouse main olfactory epithelium datasets are available under accession GSE240128; and the GAGE-seq mouse cortex and human bone marrow CD34+ datasets are available under accession GSE238001. The source code for scHiR is publicly available on GitHub at https://github.com/CSUBioGroup/scHiR.

## Acknowledgments

This work was supported by the Fundamental Research Funds for the Central Universities of Central South University and the National Natural Science Foundation of China (Grant Nos. 62225209 and 62320106009).

## Author contributions

Y.Z., R.Z. and M.L. conceived and designed the study. Y.Z. developed the computational framework and implemented the software. Y.Z. and P.J. performed the computational experiments and data analysis. M.C.H., L.T. and Z.F. contributed to the biological interpretation and validation of the results. Y.Z. prepared the figures and wrote the original manuscript. R.Z. and M.L. supervised the research. All authors reviewed and edited the manuscript and approved the final version.

